# Integration of Transplanted Interneurons Over a New Period of Ocular Dominance Plasticity in Adult Visual Cortex

**DOI:** 10.1101/2024.12.27.630358

**Authors:** Benjamin Rakela, Jennifer Sun, Philine Marchetta, Arturo Alvarez-Buylla, Andrea Hasenstaub, Michael Stryker

## Abstract

Cortical interneurons play an important role in mediating the juvenile critical period for ocular dominance plasticity in the mouse primary visual cortex. Previously, we showed that transplantation of cortical interneurons derived from the medial ganglionic eminence (MGE) opens a robust period of ocular dominance plasticity 33-35 days after transplantation into neonatal host visual cortex. The plasticity can be induced by transplanting either PV or SST MGE-derived cortical interneurons; it requires transplanted interneurons to express the vesicular GABAergic transporter; and it is manifested by changes to the host visual circuit. Here, we show that transplantation of MGE-derived cortical interneurons into the adult host visual cortex also opens a period of ocular dominance plasticity. The transplanted interneurons must be active to induce plasticity, and the neuronal activity and tuning of visually evoked responses in transplanted and host PV and SST interneurons are modulated by the locomotor state of the host. We also show that changes in activity over the period of plasticity induction are different between PV and SST interneurons but similar between host and transplanted interneurons of each type. The present findings demonstrate that the transplant-induced plasticity generated in adult visual cortex has many features in common with the role of these interneurons during the normal, juvenile critical period.

## Introduction

Cortical interneurons play an important role in mediating the critical period for ocular dominance plasticity in the primary visual cortex (V1). In mice, this period occurs over the 4-5th week of life (P21-P35) and facilitates binocular matching and optimizes visual acuity which results in a stable visual system that is well adapted to the animal’s environment. Loss of visual experience in either eye over this period, particularly the first 24 hours of deprivation, results in morphological and functional changes in V1 circuitry that favor the non-deprived eye (Espinosa and Stryker, 2012). The opening of this critical period requires GABAergic signaling from parvalbumin-expressing (PV) and somatostatin-expressing (SST) cortical interneurons from the medial ganglionic eminence (MGE). The closure is associated with factors that attenuate the GABAergic signaling of these cells (Hensch et al., 1998; Fagiolini and Hensch, 2000; Bavelier et al., 2010). Once closed, the manifestation of ocular dominance plasticity is limited, requiring greater intervention that has weaker effects on binocular vision (Espinosa and Stryker, 2012). Interestingly, transplantation of MGE-derived cortical interneurons has been shown to open a robust period of ocular dominance plasticity (33-35 days after transplantation, DAT) (Southwell et al., 2010; Larimer et al., 2016). The mechanism by which transplanted MGE-derived cortical interneurons induce a new critical period remains unknown.

Several developmental and functional processes of endogenous cortical interneurons are recapitulated by transplanted cortical interneurons. They migrate and integrate into the visual circuit, their numbers are refined by a period of programmed cell death, and the surviving cortical interneurons develop mature physiological traits (Southwell et al., 2012; Larimer et al., 2017; Bradshaw et al., 2018). The plasticity generated in the host circuit initiates changes within 24 hours of induction and follows an intrinsic timeline reflecting that of the donor (Southwell et al., 2010; Zheng et al., 2021). Plasticity can be induced by transplanting either PV or SST MGE-derived cortical interneurons, it requires the expression of the vesicular GABAergic transporter and is manifested by changes in the host circuit (Tang et al., 2014; Priya et al., 2019; Hoseini et al., 2019).

Here, we investigated the functional integration of both PV and SST transplanted cortical interneurons as they induce plasticity in the host circuit (peak at 34 DAT). We transplanted MGE-derived cortical interneurons carrying a genetically encoded calcium indicator (in either PV or SST interneurons) into adult host recipients (P44-55) that also expressed the calcium indicator in the respective interneuron type. This allowed us to compare how the neuronal activity of transplanted and host PV interneurons and transplanted and host SST interneurons are modulated by the locomotor state of the host, and how their tuning properties and activity levels change within 24 hours of monocular visual deprivation (MD). Our findings show that visually evoked responses in both transplanted and host PV and SST interneurons are larger during locomotion both before and during plasticity induction. Orientation and direction selectivity of the transplanted interneuron types were also similar to their respective hosts over this time. Changes in activity as a result of MD were different between PV and SST interneurons, but similar between host and transplanted interneurons of each type. Within 24 hours of MD, activity declined in the transplanted and host PV interneurons, whereas there was no such decline in the SST interneurons. The similarities between transplanted and host interneuron types indicate that transplanted interneurons of both types had integrated into the cortical circuit and were affected by both locomotion and MD similarly to the corresponding host interneurons.

## Results

### Transplant induced plasticity in adult primary visual cortex

Transplantation of MGE-derived PV and SST interneurons opens a new period of ocular dominance plasticity in the recipient visual cortex 33-35 DAT. In order to isolate this transplant-mediated critical period from the juvenile critical period for ocular dominance plasticity (P21-P35, peak ∼P28), we transplanted embryonic interneurons from the MGE in which either PV or SST interneurons expressed tdTomato and GCaMP7f into adult host recipients (P44-P55) that also expressed GCaMP7f in either PV or SST interneurons (Figure 1A). Experimental animals were subjected to 5 days of monocular visual deprivation (MD) by unilateral eyelid suture beginning at 33 DAT; control mice were treated identically except that they were not subject to any form of visual deprivation (Figure 1A). The effect of MD in experimental animals was measured by changes in the ocular dominance index (ODI) using intrinsic signal imaging (Cang et al., 2005). MD produced a potent and consistent ocular dominance shift in experimental animals, and no significant change in control mice (Figure 1B, Table 2). The effect of MD was like that in the juvenile critical period in that the plasticity consisted of a reduction in the response to the deprived contralateral eye (Figure 1C), rather than solely an increase in the response to the open ipsilateral eye as seen in adults. Both experimental and control ODIs prior to MD, as well as the OD shifts following MD (Figure 1D, Table 3), are consistent with published data for transplant-induced plasticity in both neonatal and adult hosts (Southwell et al., 2010, Davis et al., 2015, Zheng et al., 2021).

**Table 1.** Ocular Dominance Index (ODI) (Figure 1B) differences between PV and SST groups.

|  |  |  |
| --- | --- | --- |
|  | Two Sample Mann-Whitney Ranksum (alpha=0.05) |  |
|  | ODI<br>(group1 Median±STD vs. group2 Median±SSD) | P-value<br>(U) |
| Pre MD PV-group (n=5) vs<br>SSt-group (n=5) | 0.2900±0.0566 vs.<br>0.2900±0.0518 | 0.7698<br>(29.5) |
| Post MD PV-group (n=5) vs<br>SSt-group (n=5) | 0.0100±0.0918 vs.<br>-0.0600±0.01453 | 0.2143<br>(34) |
| Pre No MD PV-group (n=3) vs<br>SSt-group (n=3) | 0.3100±0.0802 vs.<br>0.2400±0.0451 | 0.3000<br>(13.5) |
| Post No MD PV-group (n=3) vs<br>SSt-group (n=3) | 0.2600±0.1079 vs.<br>0.2000±0.0529 | 0.4000<br>(13) |

**Table 2.** Ocular Dominance Index (ODI) (Figure 1B) Pre vs Post MD treatments.

| Table 2. Ocular Dominance Index (ODI) (Figure 1B) Pre vs Post MD treatments |  |  |  |  |  |  |
| --- | --- | --- | --- | --- | --- | --- |
|  | MD<br>(Transplant Recipients n=10;<br>n=5 PV-labeled and<br>n=5 SST-labeled) |  | No MD<br>(Transplant Recipients n=6;<br>n=3 PV-labeled and<br>n=3 SST-labeled) |  | MD + CNO<br>(Transplant Recipients n=4;<br>DREADD in PV and SST) |  |
|  | ODI<br>(Median±<br>STD) | P-value<br>(Wilcoxon<br>Sign rank,<br>alpha=0.05) | ODI<br>(Median±<br>STD) | P-value<br>(Wilcoxon<br>Sign rank,<br>alpha=0.05) | ODI<br>(Median±<br>STD) | P-value<br>(Wilcoxon<br>Sign rank,<br>alpha=0.05) |
| Pre | 0.2900 ±<br>0.0518 | 0.0020 (W=55) | 0.2650 ±<br>0.0707 | 0.3125 (W=12) | 0.2150 ±<br>0.0245 | 0.1250 (W=10) |
| Post<br>(5days) | -0.0200 ±<br>0.1195 |  | 0.2450±<br>0.0896 |  | 0.0800±<br>0.0753 |  |

**Table 3.** ODI Shift (Figure 1C)

| Table 3. ODI Shift (Figure 1C) |  |  |
| --- | --- | --- |
|  | one-way ANOVA F(2, 17) = 21.31, p < 0.0001 |  |
|  | ODI delta<br>(mean 1 vs mean 2) | P-value<br>ANOVA Post hoc multcompare |
| MD (mean 1, n=10) vs.<br>MD+CNO (mean 2, n=4) | -0.338 vs. -0.140 | 0.00854 |
| MD (mean 1, n=10) vs.<br>No MD (mean 2, n=6) | -0.338 vs. -0.0167 | 1.97 e-05 |
| No MD (mean 1, n=6) vs.<br>MD+CNO (mean 2, n=4) | -0.0167 vs. -0.140 | 0.154 |

**Figure 1.**
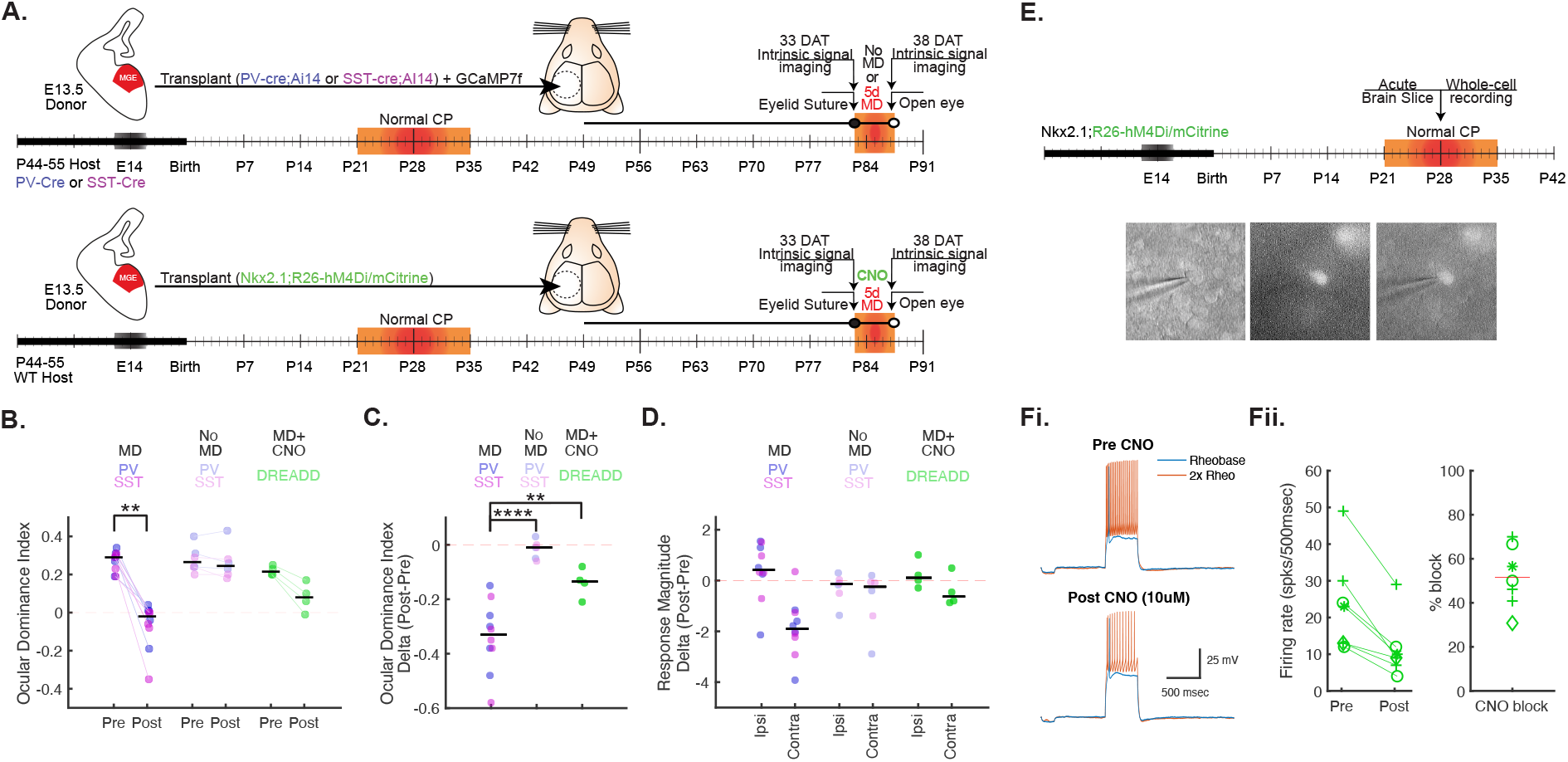
Tables 1-3.Transplant induced plasticity in adult primary visual cortex. **A)** Top. Timeline for interneuron transplantation experiments for 5 day monocular deprivation (MD) experiments and no MD experiments. The medial ganglionic eminence (MGE) containing precursors for both PV and SST interneurons (with PV or SST interneurons expressing tdTom) were transplanted into either PV or SST-cre hosts (allowing for comparison between PV interneurons or SST interneurons). Bottom. Time-line for DREADD interneuron transplantation experiments into WT hosts. The Nkx2.1-cre mouse line targets DREADD expression to both PV and SST interneurons. **B)** The Ocular Dominance Index (ODI) from Pre (33 DAT) and Post (38 DAT) timepoints is shown. Black line denotes median for the data. For each experiment involving labeled PV or SST interneuron transplants, the data are grouped together because there were no statistically significant differences within time points between the groups (Table 1). Each data point are color coded by genotype (MD group n=10, n=5 PV;Ai14 into PV-cre and n=5 SST;Ai14 into SST-cre. No MD group n=6, n=3 PV;Ai14 into PV-cre and n=3 SST;Ai14 into SST-cre. DREADD group n=4, Nkx2.1-cre;R26-hM4Di/mCitrine into WT hosts). A significant shift for the MD group was identified using a Wilcoxon Signrank test alpha=0.05. **C)** The change in ODI between Post (38 DAT) and Pre (33 DAT) time points is plotted, a one-way ANOVA revealed a significant effect, F(2, 17) = 21.31, p < 0.001, when comparing data for MD, no MD, and DREADD transplants. A post hoc comparison found that the MD group ODI shift was significantly greater when compared to the no MD or the DREADD group. **D)** The change in each eye’s response magnitude is plotted for Pre (33 DAT) and Post (38 DAT) time points, showing juvenile-like plasticity only for the MD group, where open ipsilateral eye responses increase and deprived, contralateral eye responses decrease after MD. **E)** Timeline for whole-cell current clamp experiments, where visual cortex coronal sections were made from P28 Nkx2.1;R26-hM-4Di/mCitrine mice (n=4). Brightfield, fluorescence, and merge images show recording from an mCitrine positive neuron. **Fi)** Current clamp recordings show rheobase (blue) and 2x rheobase (orange) spiking before (top) and after (bottom) 15 minutes of CNO (10uM) treatment from a representative interneuron (shown in E). **Fii)** Left. The 2x rheobase firing rate (spikes per 500msec) before and after CNO (10µM) from 7 cells from 4 mice is shown, shape denotes mouse. Right. The percent block for each cell is shown, mean block is 51.56% (red line).

Given that transplantation of either PV or SST interneurons is capable of inducing plasticity on its own (Tang et al., 2014) and that such plasticity depends on GABA release from the MGE interneurons (Espinosa and Stryker, 2012; Priya et al., 2019), we sought in another group of mice to determine whether reducing the activity of the transplanted interneurons using chemogenetics would block the plasticity induced by MD. Donor MGE-derived interneurons expressing inhibitory DREADDs were generated by crossing the Nkx2.1-cre mouse line with R26-hM4Di/mCitrine, which resulted in inhibitory DREADD receptor expression in both PV and SST interneurons (Figure 1A). These donor MGE-derived interneurons were transplanted into adult hosts as above. After baseline ODI measurements, CNO was injected IP (5mg/kg) and MD was carried out for 5 days (33-38 DAT), with CNO administration every 12 hours. While ODI shifted as a result of MD in all 4 CNO-treated animals, the shift was significantly smaller than observed in untreated animals (Figure 1B-D, Table 3), consistent with earlier findings that the activity of the transplanted interneurons is required for transplant-induced plasticity (Priya et al., 2019). To test the efficacy of DREADD-mediated inhibition in the donor Nkx2.1-cre;R26-hM4Di/mCitrine mice, acute brain sections of V1 were made and patch clamp recordings were carried out (Figure 1E-F). Washing CNO (10uM) onto acute brain slices resulted in a ∼50% reduction in firing rate in MGE-derived endogenous interneurons within 15 min. This finding indicates that CNO treatment significantly reduces the excitability of PV and SST interneurons carrying DREADD receptors but does not completely silence them.

### Activity of transplanted interneurons over the period in which they induce plasticity in adult hosts

Two-photon calcium imaging *in vivo* was applied to track the visually evoked responses of transplanted and host PV and SST interneurons expressing GCaMP7f in binocular V1 before (33 DAT) and after 24 hours of MD (34 DAT), a time shown to be critical for plasticity induction in both the juvenile critical period and in transplant-mediated plasticity (Kuhlman et al 2013, Zheng et al., 2021). At 33 DAT, imaging planes containing transplanted interneurons (either PV or SST) expressing tdTomato and GCaMP7f and host interneurons (either PV or SST) expressing GCaMP7f alone were identified. Visually evoked responses to full-field sinusoidal gratings (0.05 cy/deg) drifting at 1 cycle/sec were measured while animals were head fixed on a Styrofoam ball, where they were free to run or stay still, and locomotion was tracked using a camera (Niell and Stryker, 2010). After 24 hours of MD, the sutured contralateral eye was opened, the imaging planes from the previous day were located, and animals were again presented with the same full-field sinusoidal drifting gratings (Figure 2A). In both imaging sessions, the drifting gratings were presented 15 times at 12 directions of motion separately to each eye with presentations randomly interleaved, for a total of 360 (12 × 2 × 15) trials (Figure 2A-B). Stimuli were presented for 2 seconds and separated by inter-stimulus intervals of 3 seconds. Responses to the different stimuli were measured by deconvolution of the calcium fluorescence using Suite2p. Transplanted PV and SST interneurons expressing tdTomato as well as GCaMP7f were readily distinguished from host PV and SST interneurons. Analysis was limited to those transplanted and host PV and SST interneurons that could be matched on the subsequent imaging day and that maintained visual responsiveness (77 transplanted and 125 host PV interneurons from 5 mice, 43 transplanted and 33 host SST interneurons from 4 mice).

**Figure 2.**
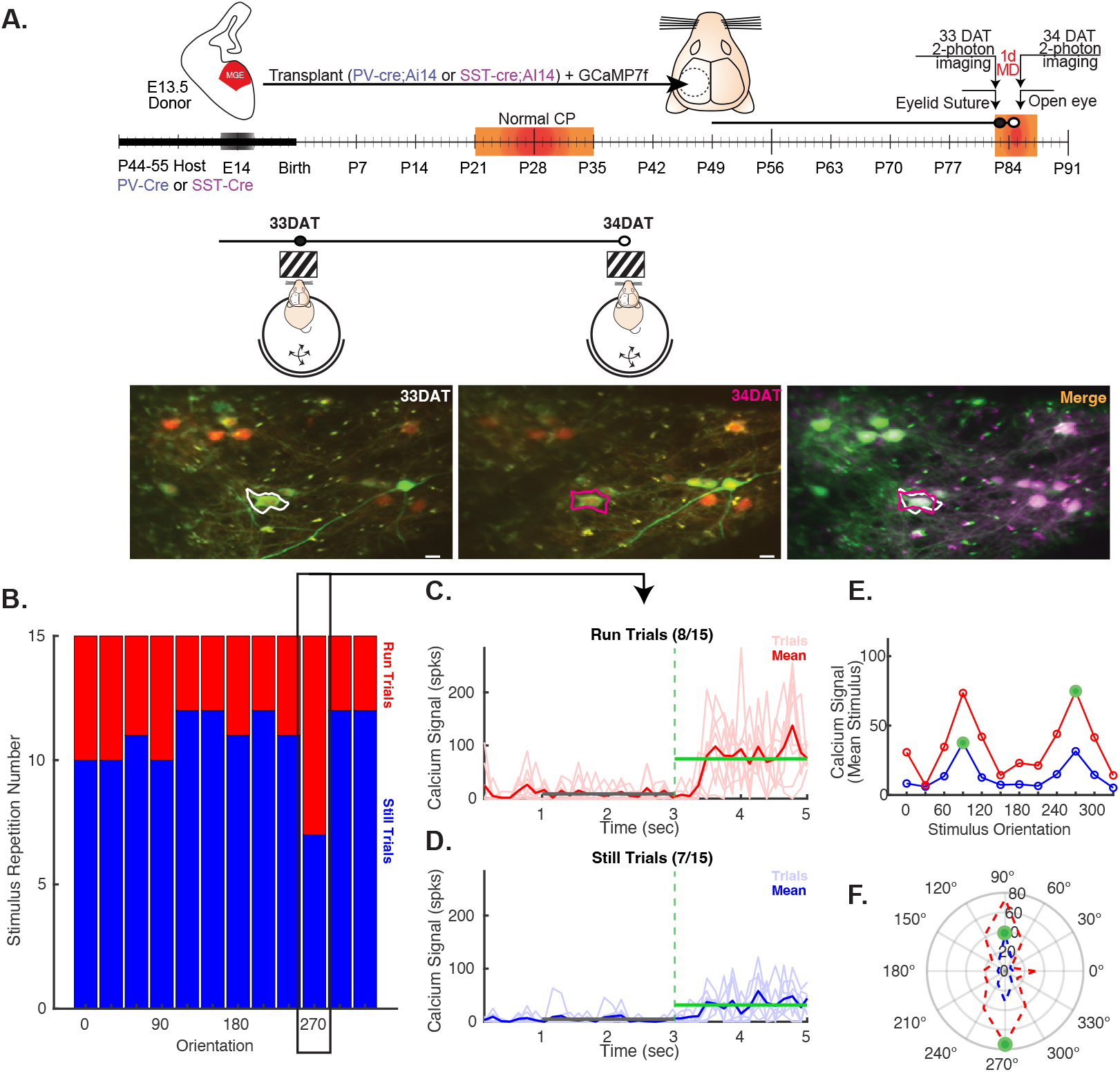
Calcium imaging of transplanted and host interneurons before and after 24 hours of MD over the transplant-mediated critical period. **A)** Timeline for interneuron transplantation experiments for 24 hours of MD experiments. The MGE containing precursors for both PV and SST interneurons (with either PV or SST interneurons expressing tdTom) were transplanted into either PV-cre or SST-cre hosts (allowing for comparison between PV interneurons or SST interneurons) along with a AAV-floxed-GCaMP7f virus. Baseline imaging was done in awake, behaving mice at a Pre-MD time point (33 DAT) and repeated for the same imaging planes after 24 hours of MD (Post time point, 34 DAT), where the same interneurons were localized. Scale bar 20 µm. **B)** The still and running trials for each orientation for contralateral eye responses from one interneuron are shown. The calcium Spks output from suite2p for 8 running **(C)** and 7 still **(D)** trials for the contralateral eye response to the 270 degree stimulus are shown. The mean for the respective trials are shown (thick red line for running (C) and thick blue line for still (D)). The mean baseline window (2 seconds-grey line) and the mean stimulus window (2 seconds-green line) are shown for the running and still trials. **E)** A tuning curve for the neuron is shown, with the largest responses identifying the preferred orientation for running trials (270 degrees) and still trials (90 degrees) represented as green dots. **F)** Polar plot for the tuning curve is shown for both running trials (red) and still trials (blue), the response magnitude is shown by the diameter of the plot and the preferred orientations are denoted by the green dots.

### Effect of locomotion on visually evoked responses during transplant-mediated critical period

In the mouse’s primary visual cortex, locomotion causes a change in cortical state that increases neuronal responses (Niell and Stryker, 2010; Dadarlat and Stryker, 2017). We measured the peak response of each neuron separately for the two eyes and for still and running trails (Figure 2 C-F). To study the effect of locomotion on transplanted and host PV and SST interneurons we computed a running index as the difference between the peak response at the preferred orientation during running and still conditions divided by their sum. Cells with a running index greater than 0 had larger responses during running trials than in still trials. Visually evoked responses through either eye were similarly increased by locomotion in both transplanted and host PV interneurons (Figure 3; Tables 3-4). This locomotion-induced increase in responsiveness remained present in both cell types through either eye after 24-hr MD. Similarly, visually evoked responses in both transplanted and host SST interneurons were also increased by locomotion before and during plasticity induction (Figure 3; Tables 3-4). The increases in the running index were independent of activity levels (Figure 3 Supplement). These data indicate that visual responses through both ipsilateral and contralateral eyes are enhanced by locomotion, and they confirm that the transplanted PV and SST interneurons are consistently integrated into and modulated by the state of the host network while they are inducing plasticity.

**Figure 3.**
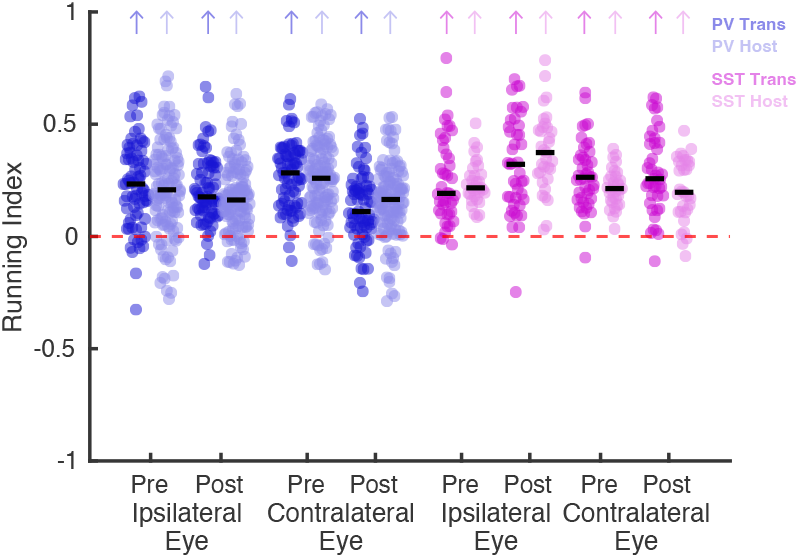
Tables 4-5. Locomotion increases the responses of interneurons to their preferred orientation. The running index (RI) for transplanted PV (dark blue), host PV (light blue), transplanted SST (dark pink) and host SST (light pink) interneurons are shown before (pre-33 DAT) and after 24 hours of monocular deprivation (post-34 DAT) for responses from the ipsilateral and contralateral eyes. Each dot represents one cell. Black line denotes median for the data. The running index was calculated from the mean response to preferred orientation for each interneuron over still and running trials (shown as the green line in Figure 2C,D for a representative neuron). Colored arrows (matching the respective cell type) indicate that the RI was greater than zero for that group, as determined by a one-sample Wilcoxon Signed-Rank Test vs 0 (alpha=0.05) (Table 4).

To quantify the effect of locomotion on the tuning of transplanted and host PV and SST interneurons we computed orientation selectivity indices (OSI) (Figure 4A-C, Table 6-7) and direction selectivity indices (DSI) (Figure 4D-F, Table 9-10) over still and running trails. Orientation (OSI) and direction selectivity (DSI) were assessed separately for the two eyes and for the running and still states by comparing the peak response of a neuron to its preferred orientation to that of the two orthogonal orientations (for OSI) or to the opposite direction of motion (for DSI). Sharply selective cells with a high OSI responded to only a few orientations of the visual stimulus. Broadly selective cells with a low OSI responded to many orientations through that eye. In some but not all conditions, transplanted interneurons, which were younger than the host neurons, were more selective (Figure 4A,B,D,E, Tables 7 and 10), consistent with the reported decline in interneuron selectivity during normal development and as transplanted interneurons mature (Kuhlman et al., 2011; Figueroa Velez et al., 2017). We then compared how these indices were affected by the locomotor state of the host by computing a running index for OSI and DSI over running and still trials for each neuron. As before, cells with an index greater than 0 had larger OSI and DSI values during running trials compared to still, whereas cells with an index <0 had larger OSI and DSI during still trials (OSI running index: Figure 4C, Table 8; DSI running index: Figure 4F, Table 11). Interestingly, the running index of OSI calculated for both ipsilateral and contralateral eye responses in both transplanted and host PV interneurons was greater than 0, indicating that the interneurons responded more selectively during running trials (Figure 4C, Table 8). The majority of host and transplanted SST interneurons, like PV interneurons, became more selective with higher OSIs during running (Figure 4C, Table 8). These locomotion-induced increases in selectivity found in both transplanted and host PV and SST interneurons were also present after 24hrs of MD. Direction selectivity (DSI) of host interneurons of both types was similar to that of the transplanted interneurons (Figure 4D-E, Table 9-10), and increased with locomotion in most cases (Figure 4F, Table 11). These data indicate the transplanted PV and SST interneurons are similarly modulated by locomotor states as the host interneurons, offering more direct evidence of their integration with the host circuit.

**Table 4.** Running Index for Peak Response (Figure 3)

| Table 4. Running Index for Peak Response (Figure 3) |  |  |
| --- | --- | --- |
|  | One-sample Wilcoxon Signed-Rank Test (vs. 0) (alpha=0.025) |  |
| | RI<br>Median ± STD | P-value (W) |
| Ipsi-eye PV trans Pre (n=77) | 0.2336 ± 0.1895 | 1.9481e-12 (2869) |
| Ipsi-eye PV host Pre (n=125) | 0.2076 ± 0.2082 | 7.8076e-17 (7287) |
| Ipsi-eye PV trans Post (n=77) | 0.1764 ± 0.1475 | 5.0921e-14 (2967) |
| Ipsi-eye PV host Post (n=125) | 0.1625 ± 0.1618 | 1.1958e-18 (7484) |
| Contra-eye PV trans Pre (n=77) | 0.2831 ± 0.1462 | 2.1731e-14 (2989) |
| Contra-eye PV host Pre (n=125) | 0.2592 ± 0.1708 | 6.1752e-21 (7718) |
| Contra-eye PV trans Post (n=77) | 0.1108 ± 0.1613 | 5.09959e-09 (2630) |
| Contra-eye PV host Post (n=125) | 0.1647 ± 0.1625 | 7.5731e-16 (7175) |
| Ipsi-eye SST trans Pre (n=43) | 0.1919 ± 0.1915 | 9.4988e-09 (939) |
| Ipsi-eye SST host Pre (n=33) | 0.2163 ± 0.0951 | 2.8224e-07 (561) |
| Ipsi-eye SST trans Post (n=43) | 0.3214 ± 0.2125 | 2.1753e-08 (927) |
| Ipsi-eye SST host Post (n=33) | 0.3736 ± 0.1733 | 2.8224e-07 (561) |
| Contra-eye SST trans Pre (n=43) | 0.2636 ± 0.1488 | 6.6851e-09 (944) |
| Contra-eye SST host Pre (n=33) | 0.2132 ± 0.0822 | 2.8223e-07 (561) |
| Contra-eye SST trans Post (n=43) | 0.2571 ± 0.1797 | 8.8569e-09 (940) |
| Contra-eye SST host Post (n=33) | 0.1968 ± 0.1395 | 8.3984e-07 (549) |

**Table 5.** Transplant vs. Host Running Index for Peak Response (Figure 3)

| Table 5. Transplant vs. Host Running Index for Peak Response (Figure 3) |  |  |
| --- | --- | --- |
|  | Two Sample Mann-Whitney Ranksum (alpha=0.025) |  |
| | RI<br>(Trans Median±STD vs Host Median±SSD) | P-value (U) |
| Ipsi PV trans (n=77) vs host (n=125) Pre | 0.2336 ± 0.1895 vs. 0.2076 ± 0.2082 | p=0.6219 (12488) |
| Ipsi PV trans (n=77) vs host (n=125) Post | 0.1764 ± 0.1475 vs. 0.1625 ± 0.1618 | p=0.1150 (12051) |
| Contra PV trans (n=77) vs host (n=125) Pre | 0.2831 ± 0.1462 vs. 0.2592 ± 0.1708 | p=0.5470 (12444) |
| Contra PV trans (n=77) vs host (n=125) Post | 0.1108 ± 0.1613 vs. 0.1647 ± 0.1625 | p=0.1513 (13267) |
| Ipsi SST trans (n=43) vs host (n=33) Pre | 0.1919 ± 0.1915 vs. 0.2163 ± 0.0951 | p=0.5715 (1325) |
| Ipsi SST trans (n=43) vs host (n=33) Post | 0.3214 ± 0.2125 vs. 0.3736 ± 0.1733 | p=0.2363 (1384) |
| Contra SST trans (n=43) vs host (n=33) Pre | 0.2636 ± 0.1488 vs. 0.2132 ± 0.0822 | p=0.0621 (1092) |
| Contra SST trans (n=43) vs host (n=33) Post | 0.2571 ± 0.1797 vs. 0.1968 ± 0.1395 | p=0.0636 (1093) |

**Table 6.** Orientation Selectivity Index (Figure 4 A-B)

| Table 6. Orientation Selectivity Index (Figure 4 A-B) |  |  |  |  |  |  |  |  |
| --- | --- | --- | --- | --- | --- | --- | --- | --- |
|  | PV- transplanted inter-<br>neuron |  | PV- Host<br>interneuron |  | SST- transplanted<br>interneuron |  | SST- Host<br>interneuron |  |
|  | Ipsi-eye<br>(Median±<br>STD) | Contra-eye<br>(Median ±<br>STD) | Ipsi-eye<br>(Median±<br>STD) | Contra-eye<br>(Median±<br>STD) | Ipsi-eye (Me-<br>dian±<br>STD) | Contra-eye<br>(Median±<br>STD) | Ipsi-eye<br>(Median±<br>STD) | Contra-eye<br>(Median ±<br>STD) |
| Still-Pre | 0.1813±<br>0.0694 | 0.1755±<br>0.07340 | 0.1590±<br>0.0707 | 0.1712±<br>0.0857 | 0.1883±<br>0.0776 | 0.2200±<br>0.1324 | 0.1428±<br>0.0485 | 0.1702±<br>0.0804 |
| Still-Post | 0.1863±<br>0.0894 | 0.1863±<br>0.1092 | 0.1408±<br>0.0815 | 0.1689±<br>0.0867 | 0.1623±<br>0.0967 | 0.2219±<br>0.1196 | 0.1312±<br>0.0618 | 0.1569±<br>0.1094 |
| Run-Pre | 0.3059±<br>0.1500 | 0.2874±<br>0.1214 | 0.3198±<br>0.1331 | 0.2677±<br>0.1180 | 0.2406±<br>0.1162 | 0.2336±<br>0.1270 | 0.2691±<br>0.1147 | 0.2046±<br>0.1379 |
| Run-Post | 0.2289±<br>0.1386 | 0.2097±<br>0.1206 | 0.2387±<br>0.1141 | 0.2278±<br>0.1230 | 0.3044±<br>0.1602 | 0.2736±<br>0.1678 | 0.3921±<br>0.1869 | 0.2615±<br>0.1371 |

**Table 7.** Transplant vs. Host Orientation Selectivity Index (Figure 4 A-B)

| Table 7. Transplant vs. Host Orientation Selectivity Index (Figure 4 A-B) |  |  |  |
| --- | --- | --- | --- |
|  |  | Mann-Whitney Ranksum (alpha=0.05) |  |
|  |  | OSI<br>(Trans Median±STD vs<br>Host Median±STD) | P-value (U) |
| Still | Ipsi PV trans (n=77) vs host Pre (n=125) | 0.1813±0.0694 vs. 0.1590±0.0707 | p=0.0097<br>(11644) |
|  | Ipsi PV trans (n=77) vs host Post (n=125) | 0.1863±0.0894 vs. 0.1408±0.0815 | p=0.0187<br>(11738) |
|  | Contra PV trans (n=77) vs host Pre (n=125) | 0.1755±0.0740 vs. 0.1712±0.0857 | p=0.4557<br>(12386) |
|  | Contra PV trans (n=77) vs host Post (n=125) | 0.1863±0.1092 vs. 0.1689±0.0867 | p=0.1464<br>(12101) |
|  | Ipsi SST trans (n=43) vs host Pre (n=33) | 0.1883±0.0776 vs. 0.1428±0.0485 | p=0.0003<br>(924) |
|  | Ipsi SST trans (n=43) vs host Post (n=33) | 0.1623±0.0967 vs. 0.1312±0.0618 | p=0.0636<br>(1093) |
|  | Contra SST trans (n=43) vs host Pre (n=33) | 0.2200±0.1324 vs. 0.1702±0.0804 | p=0.0085<br>(1019) |
|  | Contra SST trans (n=43) vs host Post (n=33) | 0.2219±0.1196 vs. 0.1569±0.1094 | p=0.0936<br>(1110) |
| Run | Ipsi PV trans (n=72) vs host Pre (n=105) | 0.3059±0.1500 vs. 0.3198±0.1331 | p=0.4894<br>(9113) |
|  | Ipsi PV trans (n=72) vs host Post (n=105) | 0.2289±0.1386 vs. 0.2387±0.1141 | p=0.9251<br>(9377) |
|  | Contra PV trans (n=70) vs host Pre (n=100) | 0.2874±0.1214 vs. 0.2677±0.1180 | p=0.5634<br>(8367) |
|  | Contra PV trans (n=70) vs host Post (n=100) | 0.2097±0.1206 vs. 0.2278±0.1230 | p=0.8880<br>(8595) |
|  | Ipsi SST trans (n=28) vs host Pre (n=7) | 0.2406±0.1162 vs. 0.2691±0.1147 | p=0.3123<br>(151) |
|  | Ipsi SST trans (n=28) vs host Post (n=7) | 0.3044±0.1602 vs. 0.3921±0.1869 | p=0.3325<br>(150) |
|  | Contra SST trans (n=40) vs host Pre (n=32) | 0.2336±0.1270 vs. 0.2046±0.1379 | p=0.4930<br>(1107) |
|  | Contra SST trans (n=40) vs host Post (n=32) | 0.2736±0.1678 vs. 0.2615±0.1371 | p=0.8695<br>(1153) |

**Table 8.**
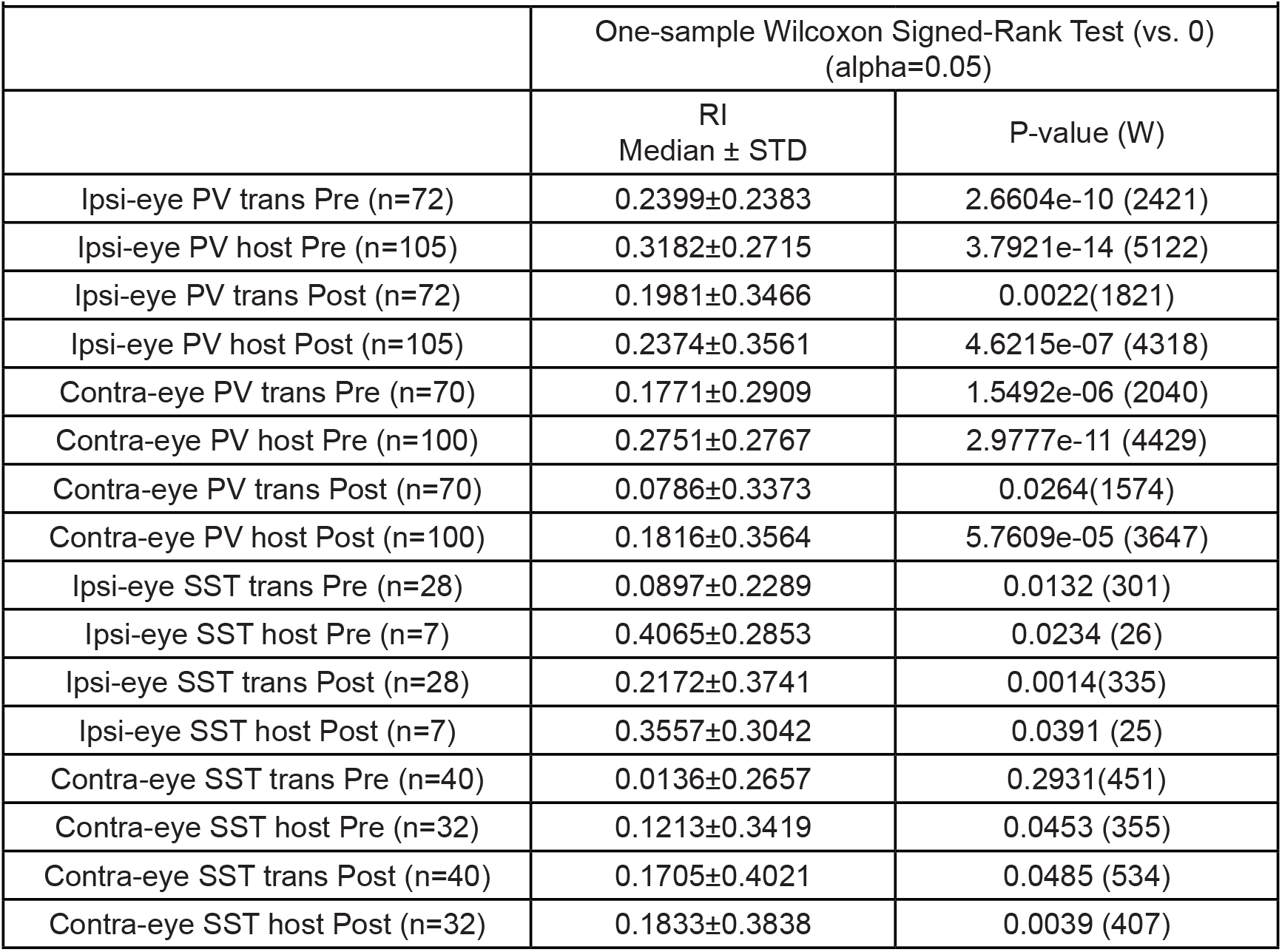
Running Index for Orientation Selectivity Index (Figure 4 C)

|  | One-sample Wilcoxon Signed-Rank Test (vs. 0)<br>(alpha=0.05) |  |
| --- | --- | --- |
|  | RI<br>Median ± STD | P-value (W) |
| Ipsi-eye PV trans Pre (n=72) | 0.2399±0.2383 | 2.6604e-10 (2421) |
| Ipsi-eye PV host Pre (n=105) | 0.3182±0.2715 | 3.7921e-14 (5122) |
| Ipsi-eye PV trans Post (n=72) | 0.1981±0.3466 | 0.0022(1821) |
| Ipsi-eye PV host Post (n=105) | 0.2374±0.3561 | 4.6215e-07 (4318) |
| Contra-eye PV trans Pre (n=70) | 0.1771±0.2909 | 1.5492e-06 (2040) |
| Contra-eye PV host Pre (n=100) | 0.2751±0.2767 | 2.9777e-11 (4429) |
| Contra-eye PV trans Post (n=70) | 0.0786±0.3373 | 0.0264(1574) |
| Contra-eye PV host Post (n=100) | 0.1816±0.3564 | 5.7609e-05 (3647) |
| Ipsi-eye SST trans Pre (n=28) | 0.0897±0.2289 | 0.0132 (301) |
| Ipsi-eye SST host Pre (n=7) | 0.4065±0.2853 | 0.0234 (26) |
| Ipsi-eye SST trans Post (n=28) | 0.2172±0.3741 | 0.0014(335) |
| Ipsi-eye SST host Post (n=7) | 0.3557±0.3042 | 0.0391 (25) |
| Contra-eye SST trans Pre (n=40) | 0.0136±0.2657 | 0.2931(451) |
| Contra-eye SST host Pre (n=32) | 0.1213±0.3419 | 0.0453 (355) |
| Contra-eye SST trans Post (n=40) | 0.1705±0.4021 | 0.0485 (534) |
| Contra-eye SST host Post (n=32) | 0.1833±0.3838 | 0.0039 (407) |

**Table 9.**
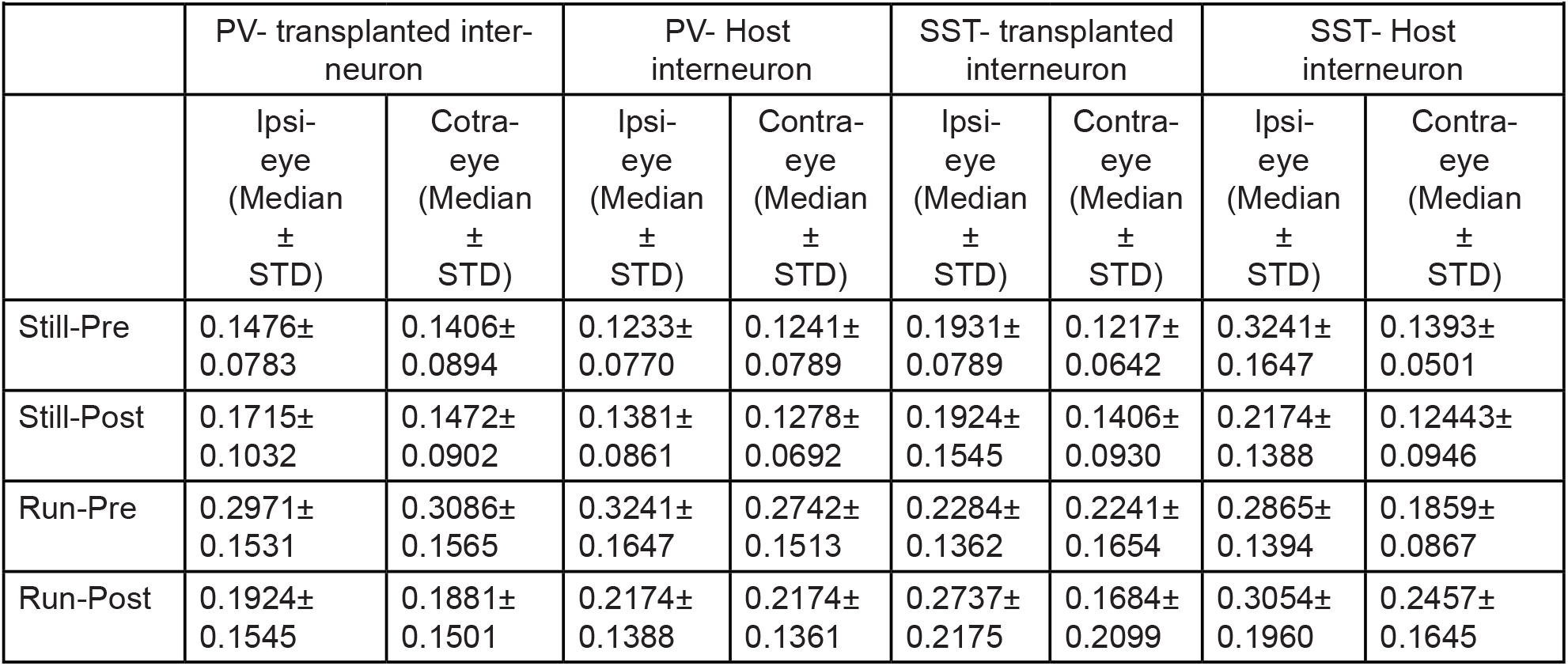
Direction Selectivity Index (Figure 4 D-E)

|  | PV- transplanted inter-<br>neuron |  | PV- Host<br>interneuron |  | SST- transplanted<br>interneuron |  | SST- Host<br>interneuron |  |
| --- | --- | --- | --- | --- | --- | --- | --- | --- |
|  | Ipsi-eye<br>(Median<br>±<br>STD) | Cotra-<br>eye<br>(Median<br>±<br>STD) | Ipsi-eye<br>(Median<br>±<br>STD) | Contra-eye<br>(Median<br>±<br>STD) | Ipsi-eye<br>(Median<br>±<br>STD) | Contra-eye<br>(Median<br>±<br>STD) | Ipsi-eye<br>(Median<br>±<br>STD) | Contra-eye<br>(Median<br>±<br>STD) |
| Still-Pre | 0.1476±<br>0.0783 | 0.1406±<br>0.0894 | 0.1233±<br>0.0770 | 0.1241±<br>0.0789 | 0.1931±<br>0.0789 | 0.1217±<br>0.0642 | 0.3241±<br>0.1647 | 0.1393±<br>0.0501 |
| Still-Post | 0.1715±<br>0.1032 | 0.1472±<br>0.0902 | 0.1381±<br>0.0861 | 0.1278±<br>0.0692 | 0.1924±<br>0.1545 | 0.1406±<br>0.0930 | 0.2174±<br>0.1388 | 0.12443±<br>0.0946 |
| Run-Pre | 0.2971±<br>0.1531 | 0.3086±<br>0.1565 | 0.3241±<br>0.1647 | 0.2742±<br>0.1513 | 0.2284±<br>0.1362 | 0.2241±<br>0.1654 | 0.2865±<br>0.1394 | 0.1859±<br>0.0867 |
| Run-Post | 0.1924±<br>0.1545 | 0.1881±<br>0.1501 | 0.2174±<br>0.1388 | 0.2174±<br>0.1361 | 0.2737±<br>0.2175 | 0.1684±<br>0.2099 | 0.3054±<br>0.1960 | 0.2457±<br>0.1645 |

**Table 10.** Transplant vs. Host Direction Selectivity Index (Figure 4 D-E)

| Table 10. Transplant vs. Host Direction Selectivity Index (Figure 4 D-E) |  |  |  |
| --- | --- | --- | --- |
|  |  | Two-sample Mann-Whitney Ranksum (alpha=0.05) |  |
|  |  | DSI<br>(Trans Median±STD vs.<br>Host Median±STD) | P-value (U) |
| Still | Ipsi PV trans (n=77) vs host Pre (n=125) | 0.1476±0.0783 vs. 0.1233±0.0770 | 0.1122<br>(12046) |
|  | Ipsi PV trans (n=77) vs host Post (n=125) | 0.1715±0.1032 vs. 0.1381±0.0861 | 0.0337<br>(11830) |
|  | Contra PV trans (n=77) vs host Pre (n=125) | 0.1406±0.0894 vs. 0.1241±0.0789 | 0.3830<br>(12335) |
|  | Contra PV trans (n=77) vs host Post (n=125) | 0.1472±0.0902 vs. 0.1278±0.0692 | 0.6484<br>(12503) |
|  | Ipsi SST trans (n=43) vs host Pre (n=33) | 0.1931±0.0789 vs. 0.1344±0.0570 | 0.0334<br>(1067) |
|  | Ipsi SST trans (n=43) vs host Post (n=33) | 0.1594±0.1103 vs. 0.1357±0.0696 | 0.0731<br>(1099) |
|  | Contra SST trans (n=43) vs host Pre (n=33) | 0.1217±0.0642 vs. 0.1393±0.0501 | 0.7933<br>(1296) |
|  | Contra SST trans (n=43) vs host Post (n=43) | 0.1406±0.0930 vs. 0.1244±0.0946 | 0.2241<br>(1154) |
| Run | Ipsi PV trans (n=77) vs host Pre (n=125) | 0.2971±0.1531 vs. 0.3241±0.1647 | 0.9460<br>(9210) |
|  | Ipsi PV trans (n=77) vs host Post (n=125) | 0.1924±0.1545 vs. 0.2174±0.1388 | 0.2512<br>(9561) |
|  | Contra PV trans (n=77) vs host Pre (n=125) | 0.3086±0.1565 vs. 0.2742±0.1513 | 0.7302<br>(9422) |
|  | Contra PV trans (n=77) vs host Post (n=125) | 0.1881±0.1501 vs. 0.2174±0.1361 | 0.1113<br>(10083) |
|  | Ipsi SST trans (n=43) vs host Pre (n=33) | 0.2284±0.1362 vs. 0.2865±0.1394 | 0.2829<br>(957) |
|  | Ipsi SST trans (n=43) vs host Post (n=33) | 0.2737±0.2175 vs. 0.3054±0.1960 | 0.3843<br>(942) |
|  | Contra SST trans (n=43) vs host Pre (n=33) | 0.2241±0.1654 vs. 0.1859±0.0867 | 0.5978<br>(1070) |
|  | Contra SST trans (n=43) vs host Post (n=43) | 0.1684±0.2099 vs. 0.2457±0.1645 | 0.3507<br>(1197) |

**Table 11.** Running Index for Direction Selectivity Index (Figure 4 F)

| Table 11. Running Index for Direction Selectivity Index (Figure 4 F) |  |  |
| --- | --- | --- |
|  | One-sample Wilcoxon Signed-Rank Test (vs. 0)<br>(alpha=0.05) |  |
|  | RI<br>Median ± STD | P-value (W) |
| Ipsi-eye PV trans Pre (n=69) | 0.3485±0.3116 | 3.0951e-09 (2180) |
| Ipsi-eye PV host Pre (n=105) | 0.3854±0.3115 | 4.9913e-15 (5204) |
| Ipsi-eye PV trans Post (n=69) | 0.08567±0.5232 | 0.3378 (1278) |
| Ipsi-eye PV host Post (n=105) | 0.2630±0.4327 | 7.7279e-06 (4135) |
| Contra-eye PV trans Pre (n=73) | 0.3630±0.3952 | 1.7324e-07 (2278) |
| Contra-eye PV host Pre (n=106) | 0.3974±0.3554 | 1.1152e-12 (5063) |
| Contra-eye PV trans Post (n=73) | 0.2382±0.4672 | 0.0089 (1782) |
| Contra-eye PV host Post (n=106) | 0.2331±0.4007 | 3.0277e-07 (4419) |
| Ipsi-eye SST trans Pre (n=37) | 0.1233±0.3355 | 0.0005 (569) |
| Ipsi-eye SST host Pre (n=27) | 0.3139±0.3337 | 0.0001 (342) |
| Ipsi-eye SST trans Post (n=37) | 0.2566±0.3350 | 0.0002 (583) |
| Ipsi-eye SST host Post (n=27) | 0.5273±0.3501 | 3.8732e-05 (354) |
| Contra-eye SST trans Pre (n=40) | 0.2685±0.3779 | 0.0004 (662) |
| Contra-eye SST host Pre (n=31) | 0.2417±0.3111 | 0.0004 (419) |
| Contra-eye SST trans Post (n=40) | 0.0480±0.4916 | 0.3071 (448) |
| Contra-eye SST host Post (n=31) | 0.2839±0.3376 | 0.0001 (437) |

**Figure 4.**
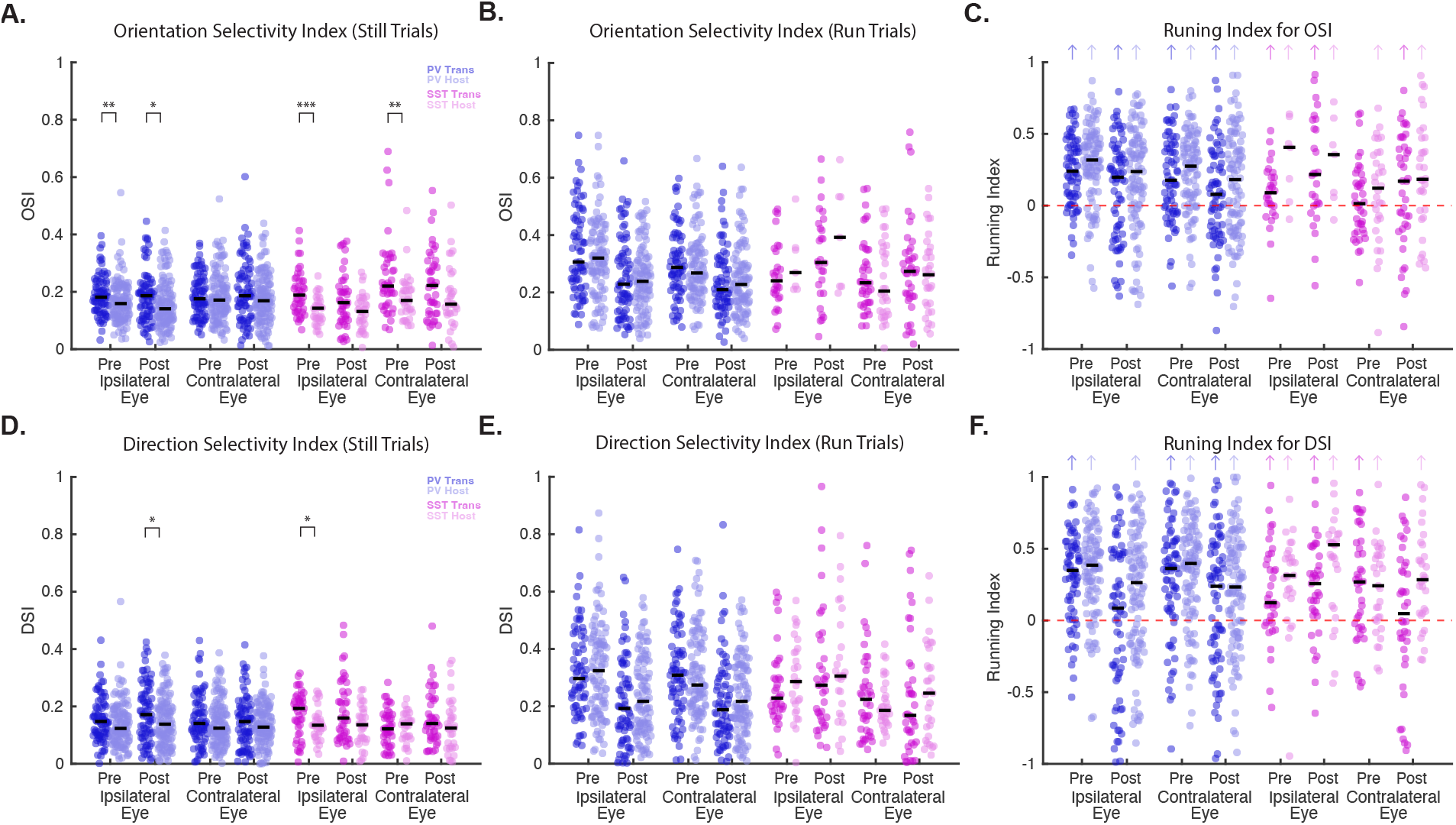
Tables 6-11. Locomotion increases the tuning of transplanted and host PV and SST interneurons. The orientation selectivity index (OSI), which was calculated from the peak response to preferred orientation and mean of the orthogonal orientations is shown for PV and SST transplanted and host interneurons from the ipsilateral and contralateral eyes for still **(A)** and running **(B)** trials. Each dot represents one cell. Black line denotes median for the data. A two-sample Wilcoxon Rank-Sum Test (alpha=0.05) found that for several transplanted vs. host interneurons OSI comparisons, the transplanted interneurons had higher OSI values in still trials, but not running trials. **C)** A running index (RI) for OSI values was calculate using the OSI values from still and running trials for each neuron. Black line denotes median for the data. Colored arrows (matching the respective cell type) indicate that the RI was greater than zero for that group, as determined by a one-sample Wilcoxon Signed-Rank Test vs 0 (alpha=0.05). **D-F)** The same analysis was done using a direction selectivity index, which was calculated from the peak response to preferred orientation and the orientation 180 degrees away for PV and SST transplanted and host interneurons from the ipsilateral and contralateral eyes for still and running trials.

### Changes in visually evoked responses following 24 hours of monocular deprivation during transplant-mediated critical period

In both the juvenile and transplant-mediated critical periods, a rapid change in PV-cell responses after 24 hours of MD has been shown to regulate plasticity; a reduction in activity of PV interneurons is thought to facilitate ocular dominance plasticity by disinhibition (Kulhman et al 2013; Zheng et al., 2021). We sought to test whether visually evoked responses in transplanted and host PV and SST interneurons were affected by 24 hours of MD. To this end, the peak response to the preferred orientation between running and still trials was measured in the same interneurons before and after 24hours of MD (Figure 2A). Visually evoked responses of transplanted PV interneurons to the deprived contralateral eye were reduced after 24 hours of MD, while responses to the non-deprived ipsilateral eye were not significantly reduced (Figure 5A,B, Table 12). The peak response to the preferred orientation in host PV interneurons was also found to be reduced after 24 hours of MD, however, this reduction was found for both ipsilateral and contralateral eye responses (Figure 5A,B, Table 12). Surprisingly, neither transplanted nor host SST interneurons exhibited reduced responses after 24 hours of MD (Figure 5C,D, Table 12). Indeed, host SST-cell responses were significantly increased by 24-hour MD, as were the responses to the ipsilateral eye of the transplanted SST interneurons (Figure 5C,D, Table 12). Differences in the activity change between PV and SST interneurons in response to 24-hour MD suggest that these interneuron types play distinct roles in mediating ocular dominance plasticity in the adult host visual cortex.

**Table 12.**
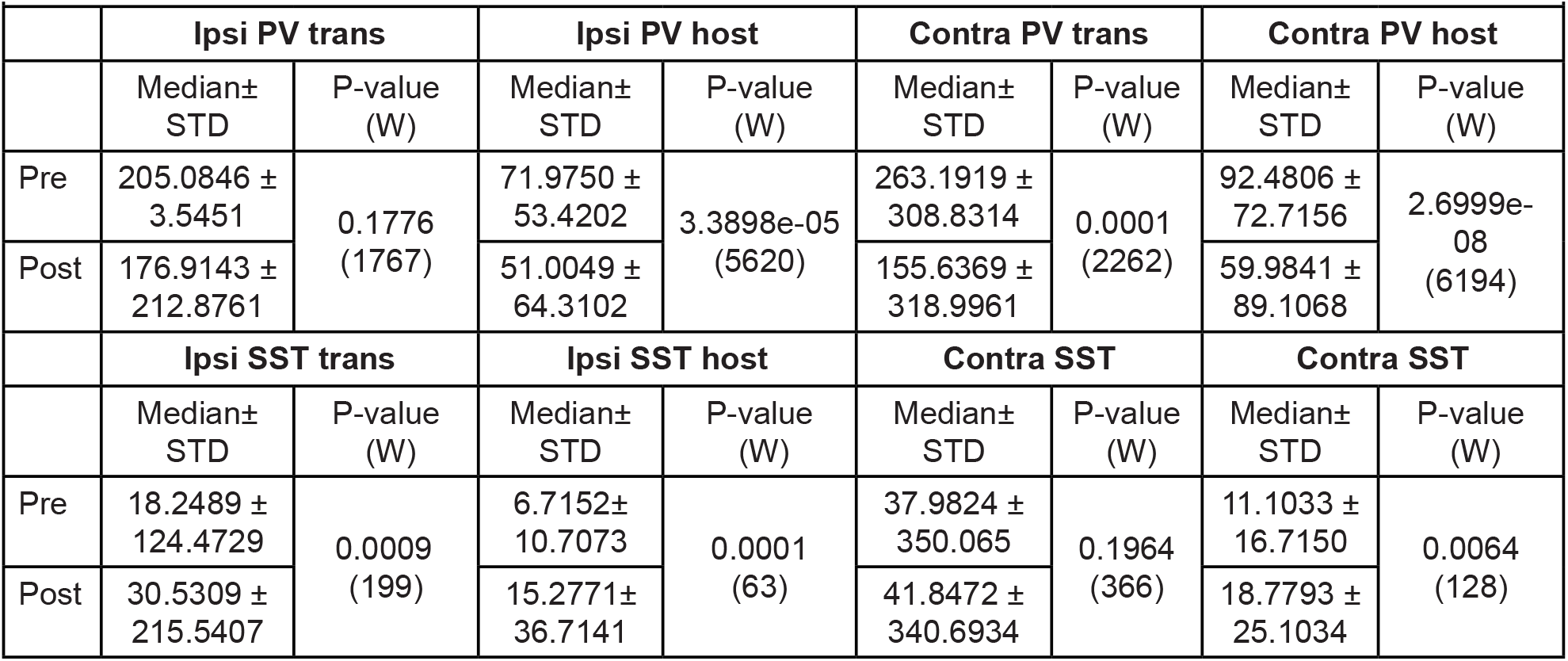
Peak Response Pre vs Post 24hrs (Figure 5 A,C) (Wilcoxon Sign rank, alpha=0.05)

|  |  |  |  |  |  |  |  |  |
| --- | --- | --- | --- | --- | --- | --- | --- | --- |
|  | Ipsi PV trans |  | Ipsi PV host |  | Contra PV trans |  | Contra PV host |  |
|  | Median±<br>STD | P-value<br>(W) | Median±<br>STD | P-value<br>(W) | Median±<br>STD | P-value<br>(W) | Median±<br>STD | P-value<br>(W) |
| Pre | 205.0846 ±<br>3.5451 | 0.1776<br>(1767) | 71.9750 ±<br>53.4202 | 3.3898e-05<br>(5620) | 263.1919 ±<br>308.8314 | 0.0001<br>(2262) | 92.4806 ±<br>72.7156 | 2.6999e-08<br>(6194) |
| Post | 176.9143 ±<br>212.8761 |  | 51.0049 ±<br>64.3102 |  | 155.6369 ±<br>318.9961 |  | 59.9841 ±<br>89.1068 |  |
|  | Ipsi SST trans |  | Ipsi SST host |  | Contra SST |  | Contra SST |  |
|  | Median±<br>STD | P-value<br>(W) | Median±<br>STD | P-value<br>(W) | Median±<br>STD | P-value<br>(W) | Median±<br>STD | P-value<br>(W) |
| Pre | 18.2489 ±<br>124.4729 | 0.0009<br>(199) | 6.7152±<br>10.7073 | 0.0001<br>(63) | 37.9824 ±<br>350.065 | 0.1964<br>(366) | 11.1033 ±<br>16.7150 | 0.0064<br>(128) |
| Post | 30.5309 ±<br>215.5407 |  | 15.2771±<br>36.7141 |  | 41.8472 ±<br>340.6934 |  | 18.7793 ±<br>25.1034 |  |

**Figure 5.**
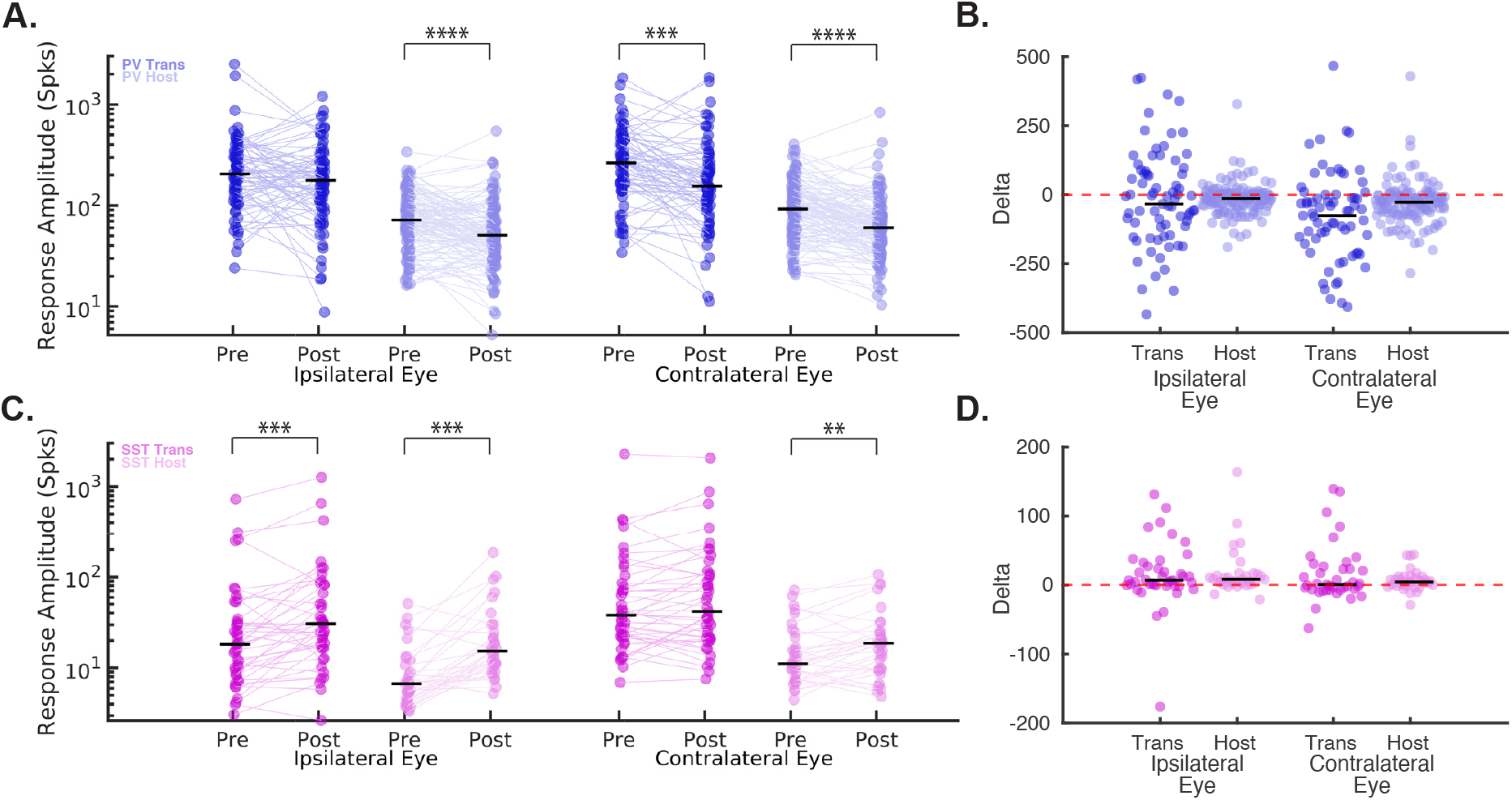
Table 12. Distinct changes in the peak responses to the preferred orientation of PV and SST transplanted and host interneurons after 24 hours of MD. **A)** Reductions in the peak response amplitude from either running or still trials to the preferred orientation were found for both transplanted and host PV interneurons for responses from the deprived, contralateral eye and were reduced for host PV interneuron for responses from the non-deprived, ipsilateral eye after 24 hours of MD (data from 77 transplanted PV interneuron pairs (dark blue) and 125 host PV interneuron pairs (light blue) are shown before (Pre-33 DAT) and after 24 hours of MD (Post-34DAT) from 5 mice, data points are paired cells color coded by genotype. Black line denotes median for the data. A two-sample Wilcoxon Signed-Rank Test (alpha=0.05) was used to compare the responses before and after MD. **B)** The change in response amplitude after 24 hours of MD (Post-Pre) for each interneuron pair from A is shown. Black line denotes median for the data. 14 transplanted PV interneurons are not shown because their delta was outside of the y-axis limits. **C)** The same analysis for 43 transplanted SST interneuron pairs, and 33 host SST interneuron pairs from 4 mice found an increase in the peak response amplitude after 24 hours of MD for most comparisons, data points are paired cells color coded by genotype. Black line denotes median for the data. A two-sample Wilcoxon Signed-Rank Test (alpha=0.05) was used to compare the responses before and after MD. **D)** The change in response amplitude after 24 hours of MD (Post-Pre) for each interneuron type from C is shown. Black line denotes median for the data. 6 transplanted SST interneurons are not shown because their delta was outside of the y-axis limits.

## Discussion

Here we confirm that transplantation of embryonic interneuron precursors from the MGE into the adult visual cortex induces a second critical period of ocular dominance plasticity, long after the end of the normal, juvenile critical period (Espinosa and Stryker, 2012). The transplanted interneurons integrate into the host circuitry so that their activity is modulated by locomotion similarly to the host interneurons. The changes produced by 24 hours of MD in transplanted PV interneurons during the transplant-induced critical period parallel those that take place in the juvenile critical period involving endogenous PV interneurons (Kuhlman et al., 2013). Namely, the activity of both transplanted and host PV cells was reduced by 24-hour MD. Interestingly, SST interneurons, both the transplanted and those in the host, were found to have increased activity over this time. This difference between PV and SST interneurons suggests that they may play distinct roles in transplant-mediated plasticity, even though each cell type has been shown to be capable of inducing plasticity on its own (Tang et al., 2014).

Reducing the activity of both transplanted PV and SST interneurons with chemogenetic approaches did impair the transplant-induced plasticity that was found after 5 days of MD. The low levels of plasticity observed in these DREADD transplants could be due to the partial block demonstrated in the brain slice recordings from mouse lines with germline DREADD expression in the PV and SST interneurons. It is possible that greater expression of DREADD receptors by viral transfection of these cells could be sufficient to completely block their activity *in vivo*. A recent paper reported that activation of inhibitory DREADDs in PV interneurons did not impair transplant-induced plasticity in adult mice (Zheng et al., 2021). This finding was interpreted as evidence that PV-interneuron activity was not involved in the plasticity phenomenon. However, this interpretation neglected the fact that, at least in cases where MGE interneuronal precursors were transplanted into the postnatal cortex in young animals, deleting the PV interneurons entirely did not impair transplant-induced plasticity (Tang et al., 2014). It was necessary to delete both SST and PV interneurons to block transplant-induced plasticity. Compromising GABA release from both PV and SST interneurons also blocked transplant-induced plasticity (Priya et al., 2019), indicating that inhibitory synaptic function is crucial for this form of plasticity.

Here, locomotion produced an increase in the orientation and direction selectivity of both transplanted and host PV and SST interneurons. Similar changes have been reported in the general population of the upper layer of V1 (Erisken et al., 2014; Dadarlat and Stryker, 2017). In several comparisons over still trials, both PV and SST transplanted interneurons were more tightly tuned than the respective host interneurons. Because the host interneurons are developmentally older than the transplanted ones, this finding is consistent with the decrease in orientation selectivity reported for PV interneurons in normal development (Kuhlman et al., 2011; Figueroa Velez et al., 2017).

Taken together, the present findings demonstrate that the plasticity generated in mature adult visual cortex by transplantation of embryonic MGE-derived precursors has many features in common with the role of these interneurons in MD during the normal, juvenile critical period.

## Methods

### Experimental Animals

All animal procedures were approved by the Institutional Animal Care and Use Committee (IACUC) at the University of California, San Francisco (UCSF) and adhered to the guidelines of the National Institutes of Health (NIH). PV-Cre;Ai14 donor mice were generated by crossing Pvalb-IRES-Cre mice with Ai14 mice (Jackson Laboratory stock numbers 017320 and 007914, respectively). Similarly, SST-Cre;Ai14 mice were produced by crossing Sst-IRES-Cre mice with Ai14 mice (Jackson Laboratory stock numbers 013044 and 007914, respectively). Mice were housed under standard conditions, including a 12-hour dark/light cycle with free access to food and water. Adult mice (P44-55) of both sexes were used for the study.

### Cell dissection and transplantation

The ventricular and subventricular zones of the medial ganglionic eminence (MGE) were dissected from E13.5 donor embryos, following the protocol described by Vogt et al. (2015). The tissue was dissociated by repeated pipetting in Leibovitz’s L-15 medium. GCaMP7f virus (pGP-AAV-syn-FLEX-jGCaMP7f-WPRE, Addgene #104492-AAV1) was added to the cell suspension and incubated for 2 hours at room temperature. AAV infection in post-mitotic cells has been shown to result in the formation of stable episomes, which remain transcriptionally active and can serve as templates for Cre-mediated recombination when Cre is expressed (Penaud-Budloo et al., 2008; Datta et al., 2024). The suspension was then concentrated via centrifugation at 800 rcf for 4 minutes. Before injection into the host brain, the cell pellet was resuspended 1:1 with the GCaMP7f virus. Adult recipients aged P44–P65 were anesthetized with isoflurane (3% for induction, 1.2–1.5% during surgery). Once the pedal reflex was absent, a craniotomy was performed over the binocular visual cortex (V1) in the left hemisphere (approximately 3 mm lateral to the midline and 1 mm anterior to lambda). The cell/virus suspension (250–350 cells/nL, 150 nL per injection for a total of ∼100,000–200,000 cells) was stereotaxically injected into three sites in the binocular V1 (50 nL per injection at 50 nL/min). Injections were made at depths of 200 and 350 μm below the pial surface using glass pipettes and a microinjection system (UMP3 UltraMicroPump, WPI). Finally, a 3-mm-diameter circular glass coverslip was affixed with cyanoacrylate to enable long-term visualization of *in vivo* neuronal calcium activity.

### Electrophysiology

To prepare acute coronal brain sections for electrophysiological recordings, animals were euthanized in compliance with approved protocols. The brain was immediately removed and placed in ice-cold dissection buffer containing (in mM): 234 sucrose, 2.5 KCl, 10 MgSO4, 1.25 NaH2PO4, 24 NaHCO3, 11 dextrose, and 0.5 CaCl2, which was oxygenated with 95% O2/5% CO2 to maintain a pH of 7.4. Coronal slices of the visual cortex (200 µm thick) were prepared using a vibratome (Precisionary Instruments) and transferred to artificial cerebrospinal fluid (ACSF). The ACSF composition was (in mM): 124 NaCl, 3 KCl, 2 MgSO4, 1.23 NaH2PO4, 26 NaHCO3, 10 dextrose, and 2 CaCl2, bubbled with 95% O2/5% CO2. Slices were incubated at 33°C for 30 minutes before being stored at room temperature. Fluorescently labeled MGE-derived interneurons (expressing mcitrine) were visualized using IR-DIC video microscopy. Whole-cell current-clamp recordings were conducted using a Multiclamp 700B amplifier (Molecular Devices) with an internal pipette solution containing (in mM): 140 K-gluconate, 2 MgCl2, 10 HEPES, 0.2 EGTA, 4 MgATP, 0.3 NaGTP, and 10 phosphocreatine (adjusted to pH 7.3, 290 mosm). After identifying the amount of current to induce one action potential (Rheobase), 2x that current was used to determine the effect of CNO. CNO (10µM) was washed on and current-clamp recordings were repeated within 15 minutes. Input resistance was monitored at the start of every recording using small hyperpolarizing pulses, and any recordings were excluded if the input resistance varied by more than 30% or if the membrane potential became unstable. Recorded signals were low-pass filtered at 2.6 kHz and digitized at 10 kHz using a 16-bit analog-to-digital converter (National Instruments). Data acquisition and analysis were performed using custom Matlab scripts.

### Intrinsic signal imaging

To elicit robust visually evoked responses, mice were administered chlorprothixene (2 mg/kg, i.m.) and maintained under light anesthesia with isoflurane (0.6–0.8% in oxygen). Core body temperature was stabilized at 37.5°C using a feedback-controlled heating system. Binocular V1 was identified using a visual stimulus spanning 20° horizontally, presented to one eye at a time while the non-tested eye was occluded with an eye shutter. The monitor was positioned 25 cm directly in front of the animal. A 2°-wide moving bar, presented at a temporal frequency of 10°/s, was generated with the Psychophysics Toolbox (Brainard, 1997; Kleiner et al., 2007) in Matlab (Mathworks) and displayed continuously (Kalatsky et al., 2003; Kaneko et al., 2008). Ocular dominance indices were determined using the same protocol, with bars oriented at 270° and 90° presented to each eye alternately and then averaged. The ocular dominance index was calculated using the formula:

### Monocular Deprivation

Monocular deprivation was induced for either 24 hours or 5 days by suturing the contralateral (right) eyelid using a 7-0 polypropylene monofilament (Ethicon) under anesthesia. The suture’s integrity was inspected daily and immediately before each visual exposure session. Animals with incompletely sealed eyelids or those whose sutures had accidentally reopened were excluded from the study. During visual exposure sessions, fullfield sinusoidal drifting gratings with 12 randomized orientations were presented to the non-deprived eye via an LED screen positioned in front of the animals, which were free to run during the sessions. On average, animals received up to 4 hours of daily visual exposure through the non-deprived eye. At the end of the deprivation period (either 24 hours or 5 days), the suture was carefully removed under anesthesia.

### Chronic, awake in vivo two-photon calcium imaging

Before baseline measurements, head-plated, transplanted mice were habituated to running or standing on a spherical treadmill. The treadmill design was adapted from Dombeck et al. (2010) and further refined by Fu et al. (2014). Visual stimuli were displayed on an LCD monitor (Dell, 30 × 40 cm, 60 Hz refresh rate, 32 cd/m^2^ mean luminance) positioned 25 cm from the mouse (−20° to +40° elevation) with gamma correction applied. Drifting sinusoidal gratings at 12 evenly spaced directions (0.05 cycles per degree, 1 Hz temporal frequency) were generated using the MATLAB Psychophysics Toolbox (Brainard, 1997; Kleiner et al., 2007) and presented in a randomized sequence. Mouse running behavior was monitored during imaging sessions with infrared illumination and a high-speed camera equipped with a 740 nm long-pass filter (Teledyne Dalsa Genie, 24.5 FPS). Running data were synchronized with calcium imaging using ball-tracking software (Neurolabware).

Imaging was performed with a resonant-galvo scanning two-photon microscope (Neurolabware, Los Angeles, CA), with data acquisition controlled by MATLAB-based Scanbox software (Neurolabware). A mode-locked Ti:sapphire laser (Coherent Chameleon Ultra II) operating at a wavelength of 920 nm provided excitation. Green fluorescence signals were collected using a 16x, 0.8 NA microscope objective with 1.7x magnification. Images were captured with a Nikon 16x water immersion objective (NA = 0.8, 3 mm working distance) in layer 2/3 of the binocular visual cortex, at depths of 150–310 μm below the cortical surface (mean ± SD: 210 ± 50 μm). Imaging was conducted at a total sampling rate of 15 Hz, distributed across two optical planes at least 30 μm apart. The same optical planes were identified across sessions using X, Y, Z coordinates and neurovascular landmarks, with MATLAB scripts employing rigid transformation analysis ensuring consistency across imaging days.

To ensure balanced visual stimulation for both eyes, randomized visual exposures were alternated between the eyes during imaging. Pneumatic eye shutters positioned in front of each eye randomly occluded one eye before each visual presentation. Each eye was exposed to 15 repetitions of randomized drifting gratings (2 seconds per stimulus) interspersed with 3-second intervals of a uniform 50% gray blank screen. During imaging sessions, head-fixed mice were free to run on a spherical treadmill (air-supported polystyrene foam ball, 20 cm diameter) while viewing the visual stimuli (Niel and Stryker, 2010).

### Data analysis

#### Calcium imaging

Cells were identified, neuropil signals subtracted, and deconvolved into putative spikes using Suite2p (Howard Hughes Medical Institute, Janelia Research Campus). Cell identities between imaging sessions were matched using custom MATLAB code (available on GitHub) and verified through visual inspection and manual correction. The Spks output from Suite2p was used for analysis. Calcium traces were processed by subtracting 70% of the respective neuropil signals and then deconvolved to estimate spike times and spike “amplitudes.” These amplitudes were proportional to the number of spikes occurring within a burst or bin. Cell selection involved an initial automated detection step using Suite2p’s cell detection algorithm, followed by manual review to ensure proper cell size and shape. Visually responsive interneurons were identified using a ranksum test. This test compared the mean response during the baseline window of each stimulus repetition to the mean response during the stimulus window across all stimulus repetitions. Analyses were performed for responses to all orientations, separately for each eye and each state.

#### Peak response

The peak response for each orientation was calculated as the best mean response during the 2-second period following stimulus onset, averaged across repetitions of that orientation for both running and still trials.

#### Running analysis

Running speed was calculated over the 2-second duration of visual stimulus presentation, with trials classified as running if the average speed exceeded 1.5 cm/s. For analyses requiring both running and still conditions, only sessions containing at least one running and one still trial for each of the 12 orientations, for both contralateral and ipsilateral eyes, were included. The running index (RI) for peak responses was derived from the peak response to the preferred orientation for each condition (mean response during the stimulus epoch for either running or still trials) for both contralateral and ipsilateral eyes. The RI was calculated using the formula:

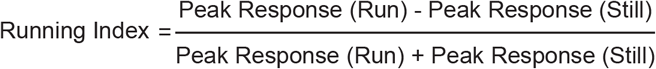

The running index (RI) for each cell’s orientation selectivity index (OSI) was derived from the OSI from still or running trials and was calculated using the formula:

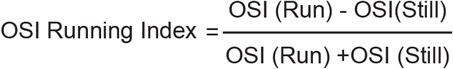

The running index (RI) for each cell’s direction selectivity index (DSI) was derived from the DSI from still or running trials and was calculated using the formula:

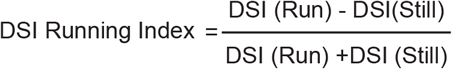

#### Statistical analysis

Statistical analyses were conducted using MATLAB (MathWorks). Statistical significance thresholds were indicated as follows: *p < 0.05, **p < 0.01, ***p < 0.001, ****p < 0.0001. Comparisons that did not reach statistical significance are not shown. The statistical tests used included: the non-parametric two sample Wilcoxon Signed-Rank Test for paired-sample comparisons, the non-parametric one sample Wilcoxon Signed-Rank Test for comparisons against zero, the non-parametric two-sample Mann-Whitney U Rank-Sum Test for unpaired comparisons, and a one-way ANOVA with post-hoc comparisons to determine whether means from different groups were significantly different. For Figure 1 ODP comparisons, the transplanted PV group and transplanted SST group were combined for 5md comparisons and noMD comparisons, respectively. A two-sample Mann-Whitney U Rank-Sum Test found no statistical difference between these groups at pre and post time points (Table 1).

## Acknowledgements

The authors would like to thank Mahmood Hoseini for his help in this study. Funding: This work was supported by NIH Grants R01-MH122478 (to A.A.B., A.R.H., and M.P.S.), F32-EY029935, K99-EY033976 (to B.R.), and R01-EY02874 (to M.P.S.). A.A.B. is also supported by a generous gift from the John G. Bowes Research Fund. A.A.B. is the Heather and Melanie Muss Endowed Chair and Professor of Neurological Surgery at UCSF. A.A.B. is Co-founder and on the Scientific Advisory Board of Neurona Therapeutics. A.R.H. is also supported by DC014101, NS116598, and EY025174, PBBR Breakthrough Fund, the Coleman Memorial Fund, and Hearing Research Inc. Author contributions: B.R. and M.P.S. conceived the experiments. B.R. performed experiments. B.R., J.S., and P.M. analyzed data. B.R. wrote the manuscript, with editing by P.M., J.S., A.A.B., A.R.H., and M.P.S.

## Supplemental Figure

**Figure 3 Supplement.**
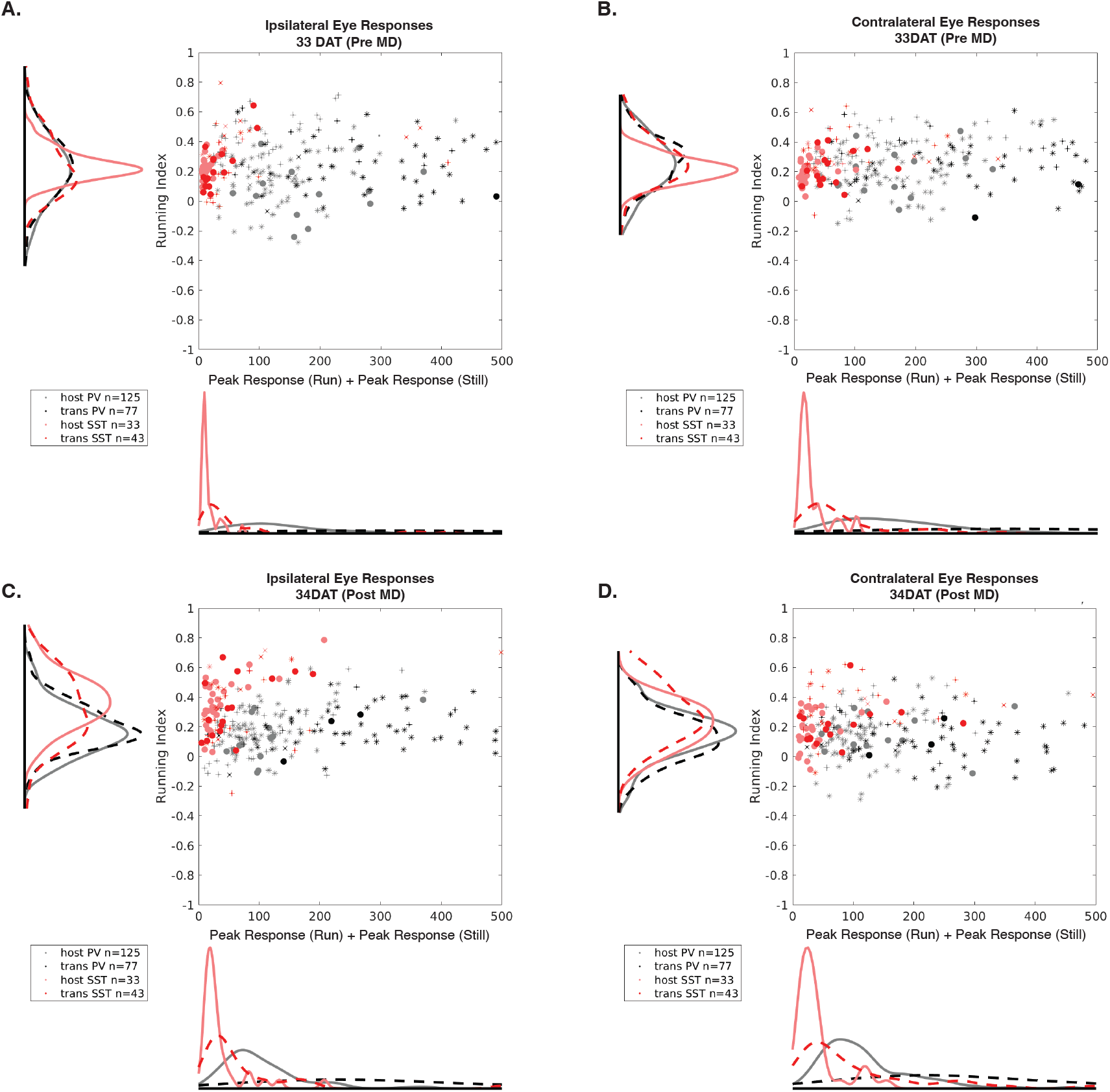
Running indices are independent of activity levels. The running index for each cell is not correlated with the amplitude of the peak response from running and still trials for ipsilateral **(A)** or contralateral **(B)** eyes before MD (33 DAT) or for ipsilateral **(C)** or contralateral **(D)** eyes after 24 hours of MD (Post-34 DAT).

## Notes

### Competing Interest Statement

Arturo Alvarez-Buylla is cofounder, serves on the scientific advisory board, and owns shares in Neurona Therapeutics.

### Summary of Updates

Revision to Acknowledgements section have been submitted.

